# Impact of tusk anomalies on the long-term foraging ecology of narwhals

**DOI:** 10.1101/2025.06.17.660143

**Authors:** Marie Louis, Alba Rey Iglesia, Jennifer Routledge, Deon de Jager, Mikkel Skovrind, Mads Peter Heide-Jørgensen, Thomas M. Kaiser, Kit M. Kovacs, Christian Lydersen, Aqqalu Rosing-Asvid, Paul Szpak, Eline D. Lorenzen

## Abstract

Male narwhals are unique in having one long, spiralled tusk, while females of the species do not have a tusk. However, a small number of individuals develop tusk anomalies, including two-tusked males or females with a tusk. In this study, we combine genetic sexing and bone collagen stable isotope (δ^13^C and δ^15^N) analysis to evaluate whether these tooth anomalies impact foraging ecology. Our analysis of individuals collected in the waters around Greenland showed no systematic impacts; nine of ten two-tusked male narwhals, and all three one-tusked female narwhals were within the normal range of known isotopic diversity from the sampled geographic regions. Two specimens with other forms of unusual dentition both showed stable isotope values outside the range of narwhals, suggesting their diet was different. Our findings underscore how DNA data retrieved from museum specimens can elucidate biological questions of interest, such as the sex of anomalous individuals. They also show how stable isotope analysis can be used to assign individuals with unknown provenance to their geographic region of origin.

## 1. Introduction

The narwhal (*Monodon monoceros*) is a High Arctic specialist found in the Atlantic sector of the Arctic. Males are known for their iconic spiralled tusk, which is an erupted left canine tooth that can grow up to 3 m in length. Males also have an embedded tusk in their right maxilla, and females have two embedded tusks in their upper maxillary bone. Neither sex has teeth in the lower jaw. The erupted tusk in males is believed to be a secondary sexual trait used to attract potential female partners, or to dissuade potential male competitors (Graham et al., 2020).

Tusk anomalies in narwhals are known to occur at low frequency; in rare cases, the embedded tusk in males grows to a full-length second tusk. Based on data from Greenland, this occurs in 0.9% of individuals. Males may also have no tusk (2.8%), and females can on occasion have a tusk (1.5%) (Garde & Heide-Jørgensen, 2022). Hay (1984) reported similar frequencies and types of narwhal tusk anomalies in Canada. There is one report of a two-tusked female; an individual collected from the Greenland Sea in 1684 that is housed at Hamburg Universität, Germany (ZMH-S-10192), which was reportedly found in association with a foetus (figure 1) (Home, 1813).

**Figure 1.**
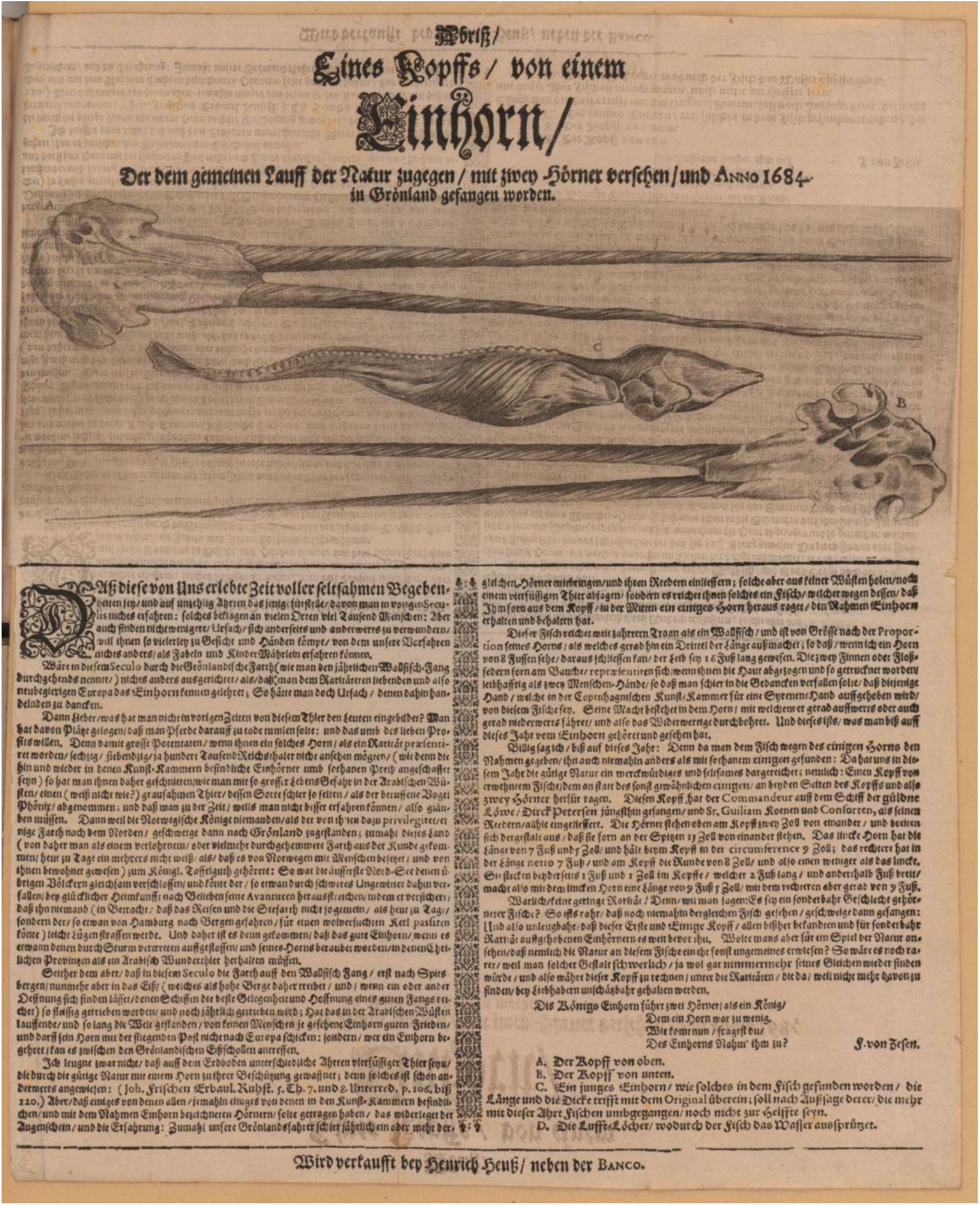
A broadsheet from 1684 detailing the find of a two-tusked female narwhal from the Greenland Sea (specimen ZMH-S-10192, see electronic supplementary material for details). The print is based on an original from the Carl-von-Ossietzky State and University Library, Hamburg.

Tooth anomalies can lead to decreased body condition in mammals (Loe et al., 2006), or to shifts in foraging ecology, as was reported in an adult, first-generation hybrid between a narwhal and a beluga (*Delphinapterus leucas*), with highly unusual dentition (Skovrind et al., 2019). This animal had teeth in both the upper maxillary and the lower jaw, as in belugas (electronic supplementary material, figure S1a) (Heide-Jørgensen & Reeves, 1993). However, several of the teeth possess longitudinal grooves and are oriented horizontally in the same manner as a narwhal tusk (and embedded tusk(s)). Bone collagen stable *δ*^13^C and *δ*^15^N isotope analysis showed elevated *δ*^13^C, indicating this hybrid animal (known colloquially as the Narluga) had a distinct diet relative to either parental species (Skovrind et al., 2019).

Narwhals live most or all of the year in areas inaccessible to humans, and much of their foraging occurs in winter (Chambault et al., 2023; Laidre & Heide-Jorgensen, 2005). Thus, biochemical markers such as stable isotopes are useful tools to gain indirect knowledge about their foraging ecology (Louis et al., 2021; Watt et al., 2013). Based on soft tissue (skin and muscle) *δ*^13^C and *δ*^15^N and on the analysis of stomach contents that provide direct data on prey consumed, narwhals feed on polar cod (*Boreogadus saida*), Arctic cod (*Arctogadus glacialis*), Greenland halibut (*Reinhardtius hippoglossoides*), capelin (*Mallotus villosus*), and squid (*Gonatus fabricii*), although the proportion of these different prey species varies in different regions (Baffin Bay, Northern Hudson Bay, East Greenland) (Garde et al., 2022; Laidre & Heide-Jorgensen, 2005; Watt et al., 2013).

In this study, we analysed a unique collection of 15 narwhals with rare tusk anomalies, to investigate the impact of dental anomalies on foraging ecology. We considered specimens with two tusks, putative females with one tusk, or individuals with other unusual tooth morphologies. Our motivation for the study was the two-tusked specimen in Hamburg (Home, 1813), presumed to be a female, and the observation of an isotopically distinct foraging ecology in the anomalous-toothed Narluga hybrid whale (Skovrind et al., 2019). By integrating genetic sexing and bone collagen stable isotope (*δ*^13^C and *δ*^15^N) analysis, we investigated the foraging ecology of the anomalous individuals – all originally collected from the waters around Greenland – relative to a panel of 84 narwhals with normal dentition.

## 2. Material and Methods

### Specimens

Our study included genetic sexing and stable isotope (*δ*^13^C and *δ*^15^N) analysis of 15 anomalous-tusked adult specimens (table 1, electronic supplementary material, table S1): ten two-tusked individuals, three one-tusked individuals believed to be females, the Narluga (a first-generation narwhal/beluga hybrid), and a narwhal with unusual dentition. Data for three specimens were sourced from the literature (Garde & Heide-Jørgensen, 2022; Rey-Iglesia et al., 2022; Skovrind et al., 2019; Vicari et al., 2022) and thus we generated novel data for 12 specimens. The specimens were originally collected in the waters around Greenland, and are housed in museums in Denmark and Germany, or at the Greenland Institute of Natural Resources (GINR).

**Table 1.** Metadata and provenance of the 15 anomalous-tusked specimens analysed. Geographic region refers to West Greenland (WG), east of Greenland (EG), or Greenland (GL, if the specific region of origin is unknown). The substrate used for DNA and stable isotope analysis (SIA) is indicated.

| Museum ID | Tusk anomaly | Locality | Region | Genetic sex | Substrate analysed (DNA/SIA) | Collection date |
| --- | --- | --- | --- | --- | --- | --- |
| M08-CN1x | Two-tusked | NA | GL | M | tusk/bone | 1800s |
| M08-CN10x | Two-tusked | Itivdlarsuk | WG | M | tusk/bone | Oct 1883 |
| M08-CN11x | Two-tusked | Upernavik | WG | M | tusk/bone | May 1884 |
| M08-CN35x | Two-tusked | NA | GL | M | tusk/bone | 1897 |
| M08-CN58 | Two-tusked | Kap York | WG | M | tusk/bone | 1921 (reg. date) |
| M08-CN76 | Two-tusked | NA | GL | M | tusk/bone | 1965 (reg. date) |
| M08-CN2x | Two-tusked | Uummannaq | WG | M | tusk/bone | 1800s |
| NA | Two-tusked | NA | GL | M | tusk/bone | NA |
| ZMH-S-10192 | Two-tusked putative female | Greenland Sea | EG | M | tusk/bone | 1684 |
| 1029 | Two-tusked | Scoresby Sound | EG | M | skin/NA | 2009 |
| 1196 | One-tusked putative female | Kitsissuarsuit | WG | F | tusk/tusk | 1996 |
| 1197 | One-tusked putative female | Kitsissuarsuit | WG | F | tusk/tusk | 1996 |
| 938 | One-tusked female | Scoresby Sound | EG | NA | (Garde & Heide-Jørgensen, 2022)/(Rey-Iglesia et al., 2022) | 2017 |
| M08-CN44 | Odd dentition/unusual teeth | NA | NA | M | (Vicari et al., 2022)/(Vicari et al., 2022) | 1963 |
| MCE1356 | Odd dentition/unusual teeth | Kitsissuarsuit | WG | M | (Skovrind et al., 2019)/(Skovrind et al., 2019) | 1986 |

Specimen ZMH-S 1019 has been presumed to be female. It was brought to Hamburg, Germany, in 1684 after being harvested from the Greenland Sea by the German whaling vessel De Goude Leeuw. A contemporary news bulletin referred to the individual as female, due to a foetus found in association with the adult animal (figure 1) (Home, 1813).

We generated DNA and stable isotope data for nine of the ten two-tusked specimens available for analysis; as we had only skin tissue for the remaining individual (1029), it was only genetically sexed (table 1). We also generated novel DNA and stable isotope data from two putative one-tusked females; data from a third individual was sourced from the literature (Rey-Iglesia et al., 2022). Both tusks have an unusual shape and colour (electronic supplementary material, text and figure S1b). We also included data sourced from the literature for two individuals with highly unusual dentition, both genetically identified as males: the Narluga (Skovrind et al., 2019) (MCE1356) and another narwhal individual (CN44). The latter had a cranial shape similar to narwhals, yet had a row of several erupted teeth in the upper maxillary (electronic supplementary material, figure S1c) (Vicari et al., 2022).

As a reference panel, we included bone collagen stable isotope *δ*^13^C and *δ*^15^N data from 84 narwhals (figure 2; electronic supplementary material, figure S2): data from 40 and 39 individuals from West and East Greenland, respectively (Louis et al., 2021), and five individuals from Svalbard, from which novel data were generated to expand the geographic range of the reference material (electronic supplementary material, table S1).

**Figure 2.**
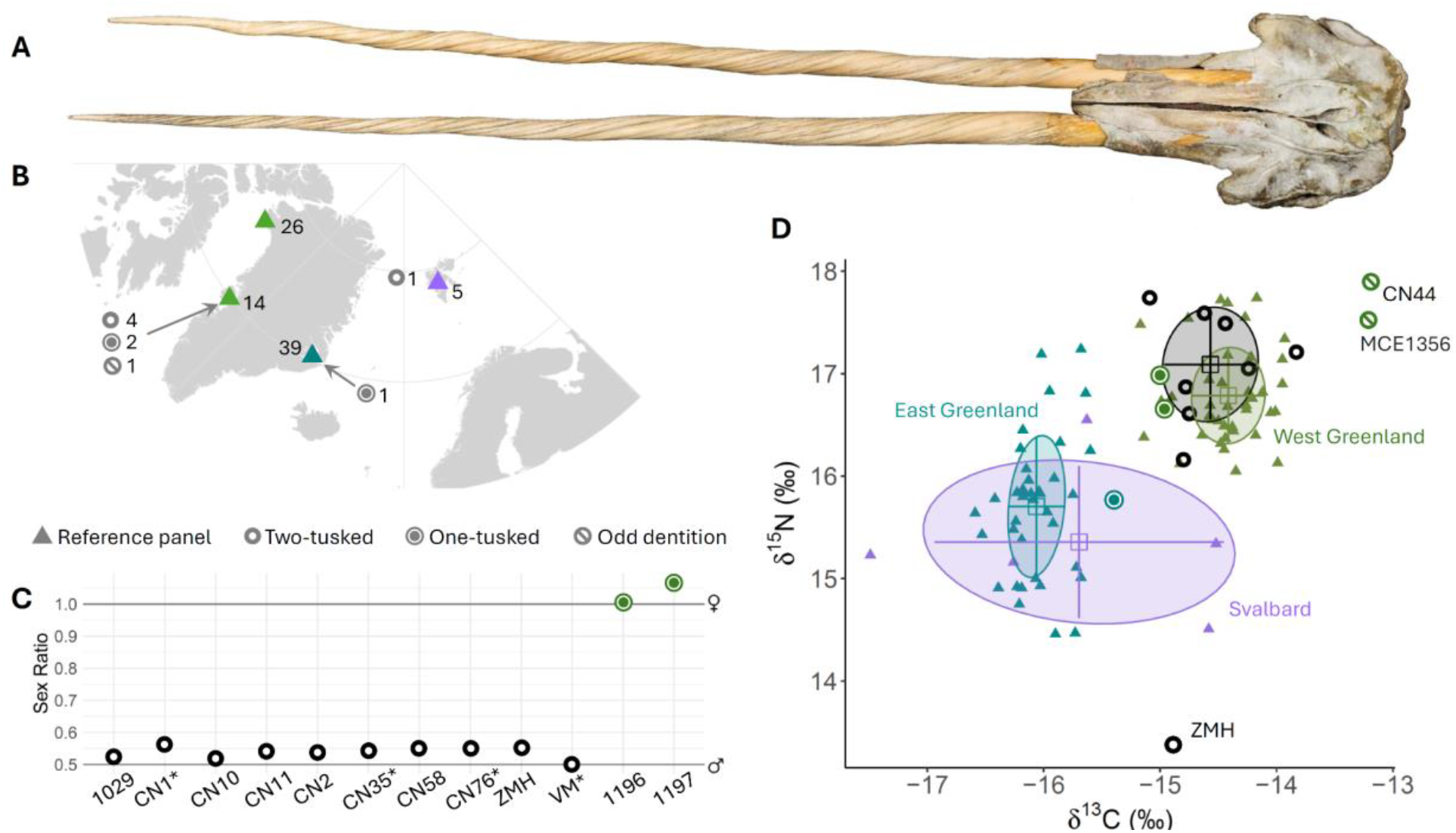
Samples and biomolecular results. (A) Photograph of specimen ZMH-S-10192, courtesy of Mentz/Centrum für Naturkunde -Universität Hamburg. (B) Map showing the sampling localities of the anomalous-tusked individuals with known geographic provenance in grey, and the 84 narwhals used as a reference panel in (D): West Greenland in green, East Greenland in teal, Svalbard in purple. (C) Genetic sexing of ten two-tusked (black) and two one-tusked (green) individuals. Genetic sex was determined by estimating the X chromosome:autosome coverage ratio (X:A ratio); ratio values ∼0.5 indicate the individual is a male, and ∼1.0 a female. * indicates the known provenance is only Greenland. (D) Bone collagen *δ*^13^C and *δ*^15^N of the 15 anomalous-tusked individuals analysed and the reference panel. Shaded ovals indicate Bayesian standard ellipse areas for the individual groups (SEA_B_). The SEA_B_ of the double-tusked individuals (black) was estimated based on the eight individuals that were sampled in or assigned to West Greenland. Mean (square) and SD (error bars) are shown. Individuals discussed in the text are indicated.

### Laboratory analyses

#### DNA

We drilled 50-70 mg of dentine powder from nine two-tusked narwhals and two putative one-tusked females (table 1). DNA extractions, library builds, and sequencing are detailed in the electronic supplementary material.

#### Stable isotopes

We sampled 300 mg of bone powder from the skulls of nine two-tusked narwhals, and tusk powder from two putative one-tusked females, as no bone was available (table 1). We followed the approach described in Louis *et al*. (2021), detailed in the electronic supplementary material. To analyse bone and dentine (tusk) data together, dentine isotopic values need to be translated into bone-equivalents using a correction factor, and we thus applied a correction factor (Rey-Iglesia et al., 2022) to the *δ*^13^Cand *δ*^15^N values obtained from the two tusks for which we did not have accompanying bone. We adjusted the *δ*^13^C values to correct for change in atmospheric and oceanic dissolved inorganic carbon, which has occurred since the late 19th century due to industrialization (the Suess Effect; (Keeling et al., 1979; Quay et al., 1992)). If the collection date was not available, the registration date into the museum collections was used to correct for the Suess effect (table 1).

### Data analyses

#### Genetic sexing

We used the SEXY pipeline to identify the sex of our sampled individuals (Cabrera et al., 2022). Briefly, we determined sex by estimating the X chromosome:autosome coverage ratio (X:A ratio) after mapping the sequencing reads to a narwhal reference genome (GCF_005190385.1). Specimens with X:A ratio < 0.7 were determined as males, X:A ratio > 0.8 as females (see details in the electronic supplementary material).

#### Stable isotopes

We ran all of our statistical analyses in R v.3.6.1 (R Core Team, 2019). We statistically compared the two-tusked narwhals – apart from specimen ZMH-S-10192, which was sampled in the Greenland Sea – with the reference data set for narwhals collected in the same region (West Greenland). We also compared the two-tusked narwhals with the reference data separating males and females, see electronic supplementary material for details and results. For the three one-tusked females, the Narluga, and the narwhal with unusual erupted teeth, sample sizes were too low to run statistical analyses, and thus we evaluated the results visually.

Our data satisfied the assumption of normality and homogeneity of variance for the different levels of subdivision. To test for differences in foraging ecology between the eight two-tusked narwhals (four from West Greenland and four with unknown locality in Greenland, but which grouped with the west) and the reference data from narwhals from West Greenland (n=40), we compared their *δ*^13^C and *δ*^15^N using Student’s t-tests.

Using isotopic niche as a proxy for ecological niche, we compared the isotopic niches of the eight two-tusked narwhals and the West Greenland reference data, using Bayesian multivariate ellipse-based metrics implemented in the packages SIBER and rjags (Bearhop et al., 2004; Jackson et al., 2011; Newsome et al., 2007; Plummer et al., 2016). We calculated standard ellipse areas corrected for sample size (SEA_C_), and Bayesian standard ellipses (SEA_B_) for each group. We estimated SEA_B_ using 10^5^ posterior draws, a burnin of 10^3^ and a thinning of 10, and used SEA_B_ to test for differences in niche width among groups (i.e., the proportion (p) of draws of the posterior distribution of the SEA_B_ in which the area of one group was smaller than the other). We evaluated isotopic niche similarity between two groups as the proportion (%) of the non-overlapping area of the maximum likelihood (ML) fitted ellipses of the two. We generated SEA_B_ for the five Svalbard individuals as well; the individuals were not sexed and were thus pooled. We did not run any statistics on this region due to the low sample size.

## 3. Results

The ten two-tusked individuals were all identified as male, with X:A ratios between 0.50 and 0.56 (figure 2C, electronic supplementary material, table S2). This included specimen ZMH-S-10192, which was assumed for the past 340 years to be a female (figure 1). The two other putative one-tusked females analysed were confirmed to be females, with X:A ratios between 1.0 and 1.07.

We did not have specific locality information for all individuals analysed, and four of the two-tusked individuals were known only to originate from ‘Greenland’ (table 1). Narwhals in West and East Greenland are isotopically differentiated (Louis et al., 2021). The four two-tusked narwhals which were known *a priori* to originate from West Greenland grouped with the reference dataset from this region (figure 2D). The four specimens with only ‘Greenland’ as known provenance similarly grouped with the West Greenland narwhals, we thus assigned these individuals to an origin in West Greenland, and included the total of eight samples in the statistical analysis. We did not find any significant difference in *δ*^13^C and *δ*^15^N between the two-tusked individuals and the reference panel (t=-1.09, df = 8.81, P = 0.30 for *δ*^13^C; t= 1.45, df = 9.22, P = 0.18 for *δ*^15^N, figure 2D). Their isotopic niche (SEA_B_=0.75 ‰^2^) was not significantly different (proportion p=0.86) from the reference panel (SEA_B_=0.46 ‰^2^, figure 2D). Ellipse overlap between the two isotopic niches was 35%.

Due to the absence of bone collagen δ^13^C and δ^15^N data from animals in Svalbard, we generated novel data from five animals, including three stranded animals, that were opportunistically sampled in the region (electronic supplementary material, table S1). To ascertain if narwhals from East Greenland and from Svalbard are isotopically differentiated, we generated Bayesian standard ellipses for each of the three sampled reference populations (also incl. West Greenland, figure 2D). Although based on a limited sample size, the range of δ^15^N observed in Svalbard was within the range observed in East Greenland.

The δ^13^C values of the Svalbard individuals had a wide range, overlapping with both West and East Greenland, which may reflect some of the samples being derived from stranded specimens, and thus perhaps originating from other regions. The isotopically derived ecological niche of the Svalbard individuals encompassed most of the niche of East Greenland narwhals (figure 2D).

The bone collagen *δ*^13^C value for specimen ZMH-S-10192 was within the range of values observed in narwhals from Svalbard (figure 2D). However, the individual’s *δ*^15^N was much lower than any other sample included in our analysis. Visual inspection of the stable isotope values retrieved for the other anomalous-tusked individuals, where sample size prevented statistical analyses, indicated the two West Greenland one-tusked females had values within the range observed for the West Greenland female reference panel (electronic supplementary material, figure S2). The one-tusked female from East Greenland had *δ*^15^N values within the range observed for the East Greenland female reference panel, although the *δ*^13^C value of this individual was somewhat higher (by 0.27‰). Specimen CN44, the male narwhal with unusual teeth, had a higher *δ*^13^C value than any other narwhal analysed, similar to the Narluga (Skovrind et al., 2019).

## 4. Discussion

By combining genetic sexing and bone collagen stable isotope (*δ*^13^C and *δ*^15^N) analysis, we evaluate whether tusk anomalies in narwhals impact foraging ecology. δ^15^N provides information on the trophic level(s) at which animals are feeding, and δ^13^C provides information on foraging habitat (pelagic versus demersal) [43–46]. Due to the relatively slow turnover of bone collagen, stable isotope data represent the diet of individuals over several years to decades (Hedges et al., 2007; Wild et al., 2000). We hypothesised that tooth anomalies impact foraging ecology, as has been observed in the Narluga (Skovrind et al., 2019), albeit perhaps to varying degrees.

We compared the narwhals with tusk anomalies with a reference panel of narwhals from West and East Greenland (Louis et al., 2021), and from Svalbard. Four of the two-tusked specimens had known provenance in West Greenland (table 1), and they clearly clustered with the reference narwhals from the region (figure 2D). The other four two-tusked specimens, for which we did not have specific locality information (the museum ledgers only mention ‘Greenland’), also clustered clearly with the reference narwhals from West Greenland. This is unsurprising; the majority of the collections at the Natural History Museum of Denmark originating from Danish voyages to the Arctic during the period when the specimens were collected (table 1), are from the west coast of Greenland.

Based on this, we analysed the eight individuals together and found no significant differences in their δ^13^C or δ^15^N and the reference panel from West Greenland (figure 2D). The findings suggest having two tusks does not impact isotopically-derived resource use. However, we acknowledge that our sample size is low, and thus our findings should be interpreted with caution. We assumed isotopic stability across the 200 years covered by the West Greenland samples (table 1), implying that there has been no significant shift in stable isotope values apart from the Suess effect, which we account for in our analyses. This is supported by a comparable bone collagen δ^13^C and δ^15^N analysis of belugas; belugas sampled from their summering ground in Elwin Bay, High Arctic Canada, between 1874 and 1898, and from their West Greenland wintering grounds in the 1990s show no significant shifts in δ^13^C and δ^15^N across time (Szpak et al., 2020).

Specimen ZMH-S-10192 was caught in the Greenland Sea, between East Greenland and Svalbard (figure 2D). The specimen has been assumed to be a female, but our genetic sexing clearly showed it was a male, as all of the other double-tusked specimens (figure 2C). The isotopically-derived foraging ecology of this animal differed markedly from the reference panel, with a δ^13^C value higher than East Greenland narwhals, but similar to two of the Svalbard individuals (figure 2D). The δ^15^N value for this animal was 1.1 ‰ lower than the lowest values observed in East Greenland/Svalbard, indicating ZMH-S-10192 was likely feeding at a lower trophic level than narwhals in general. The narwhals in the East Greenland reference panel were caught between 1993 and 2007 (Louis et al., 2021), and the Svalbard individuals (for which collection dates exist) were collected between 1886 and 2014 (table 1). ZMH-S-10192 was collected in 1684 and thus differs from the other two-tusked narwhals by its much older age. However, no temporal stable isotope data are available from East Greenland. Thus, we cannot rule out that the lower δ^15^N value may be due to ecological changes over time, or shifts in δ^15^N at the base of the food web due to reconfiguration of the primary producers, which would affect values observed in top predators, even if the latter feed on the same prey or same trophic level (Sherwood et al., 2014). Thus, with the data available, we cannot further elucidate why we observe a low δ^15^N value in ZMH-S-10192.

The two one-tusked females from West Greenland had isotopic values within the range of the reference data from the region (figure 2D); the higher value of δ^13^C in the one-tusked female from East Greenland relative to the reference panel is within the precision range of the method. The lack of a systematic difference in the isotopically-derived foraging ecology of anomalous-tusked narwhals relative to normal-tusked narwhals suggests the presence of an extra tusk does not impact narwhal foraging ecology significantly. Additionally, the one-tusked female from East Greenland, which was a carcass examined by biologists, showed signs of multiple pregnancies (Garde & Heide-Jørgensen, 2022), indicating that the presence of a tusk does not interfere with female reproductive success.

The Narluga is from West Greenland (Skovrind et al., 2019), but the sampling locality of specimen CN44 is unknown (Vicari et al., 2022). The latter had elevated δ^15^N and δ^13^C – similar to values observed in the Narluga. The elevated δ^13^C may indicate a more demersal foraging ecology, as has been posited for Narluga (Hobson et al., 1995; Skovrind et al., 2019). As our stable isotope values are from bone, which has a slow turnover of years to decades, the isotopic compositions presented in this study reflect general, long-term foraging ecology (Hedges et al., 2007; Wild et al., 2000). Although based on only two specimens, our findings suggest the presence of beluga-like erupted teeth does impact foraging ecology, possibly allowing individuals to capture or handle other types of prey than other (normal) narwhals. We cannot determine prey species based on our approach, but the elevated *δ*^15^N values were in the top of the range observed in narwhals.

Our findings suggest the vast majority (12 out of 13) of the two and one-tusked anomalous individuals analysed are within the isotopic niche of normal-tusked narwhals, and thus tusk anomalies do not appear to systematically impact foraging ecology. Our analysis also illustrates the potential of using stable isotopes to assign specimens to a geographic region of origin, as we were able to do for the four two-tusked individuals of unknown provenance. Narwhal ivory has been internationally traded for hundreds of years, since the Norse trade in the Middle Age (Reeves & Heide-Jørgensen, 1994). DNA, and mitochondrial DNA in particular, has been applied to investigate the geographic origin of ivory from other species (Ruiz-Puerta et al., 2023; Star et al., 2018). However, there is no phylogeographic structuring in narwhal mitochondrial DNA (Louis et al., 2020), and our findings suggest that δ^13^C and δ^15^N analysis may be a promising avenue for investigating spatiotemporal patterns of the narwhal ivory trade.

## Data Accessibility Statement

Electronic supplementary material, table S1 for the stable isotope data. The raw sequencing files, input files and scripts will be available at the Electronic Research Data Archive at University of Copenhagen.

## Authors contributions

ML and EDL conceived the project. MPHJ, KMK, TMK, CL, ARA provided samples. ML, MS, TMK, EDL sampled the specimens. ARI, JR, DJ and PS generated the data. ML and ARI analysed the data. ML and EDL wrote the manuscript and all co-authors edited the manuscript.

## Acknowledgements

We thank Marianne Sørensen and Daniel Klingberg Johansson for help with access to the collection and metadata of the double-tusked specimens from Varde Museum and the NHMD respectively. We also thank Matthias Preuss for assistance with the resampling of the Hamburg specimen; Daniel Bein and Matthias Glaubrecht (Hamburg) as well as Christiane Caemmerer (Berlin) for valuable information as to the historical documentation of the Hamburg specimen, and we thank Sander Solnes, Lars Erik Johannessen and Daniela Kalthoff for access to specimens from the Svalbard Museum, the Natural History Museum Oslo and the Swedish Museum of Natural History, respectively. We thank Eva Garde for her input on the manuscript, Camilla Hjorth Scharff-Olsen and Lennart Schreiber for their help in the DNA lab. We also thank the Greenlandic hunters who provided the tusks.

This research was supported by the Carlsberg Foundation Distinguished Associate Professor Fellowship, grant CF16-0202 to EDL. ML was supported by a postdoctoral fellowship from the Greenland Research Council. Svalbard sample collections were supported by the Norwegian Polar Institute.

