## Supplementary material for "Impact of tusk anomalies on the long-term foraging ecology of narwhals"

Overview of the supplementary material

Supplementary figure S1

Supplementary tables S1 and S2

Supplementary text

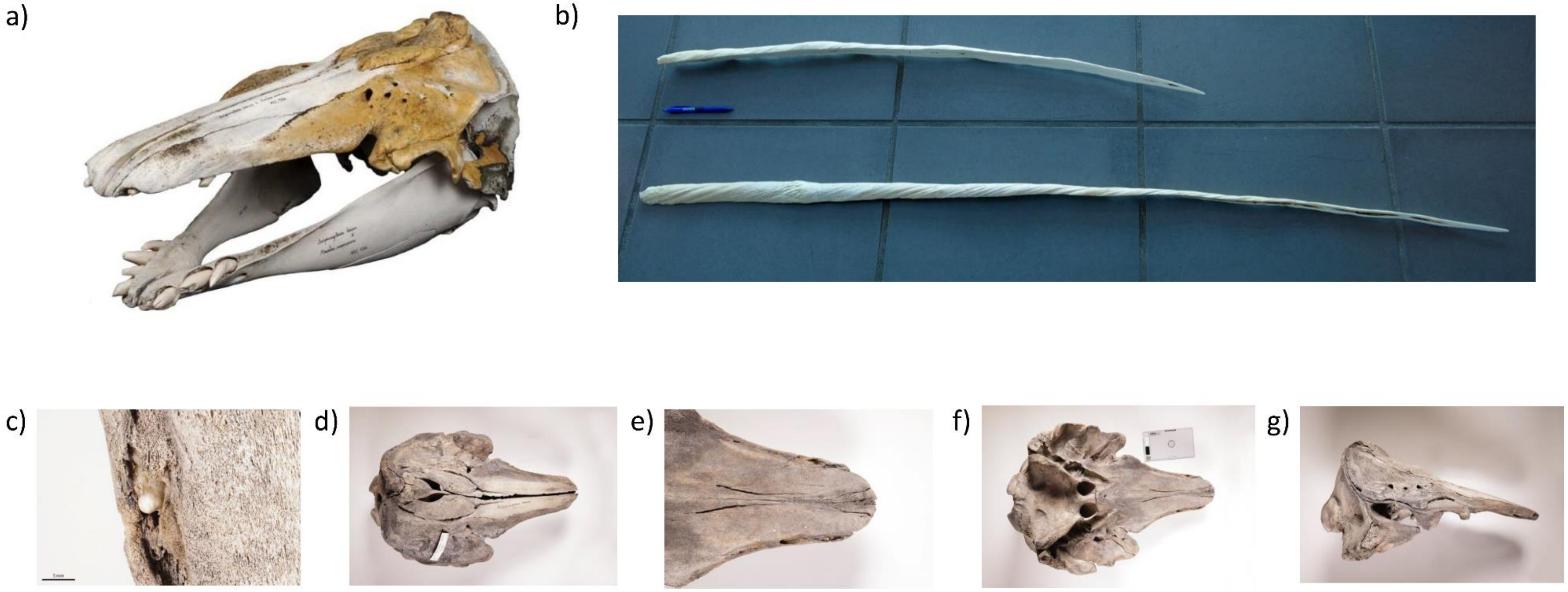

**Supplementary figure S1. Photos of four of the anomalous specimens.** a) Skull of specimen MCE1356, a first-generation hybrid between a narwhal and a beluga, also called Narluga, photo: Mikkel Høegh Post. b) The single tusks from two putative females (1197 at the top and 1196 at the bottom); both tusks have been sectioned, photo: Fernando Urgate. c-g) Five images of the anomalous narwhal specimen (M08-CN44), photos: Mikkel Høegh Post.

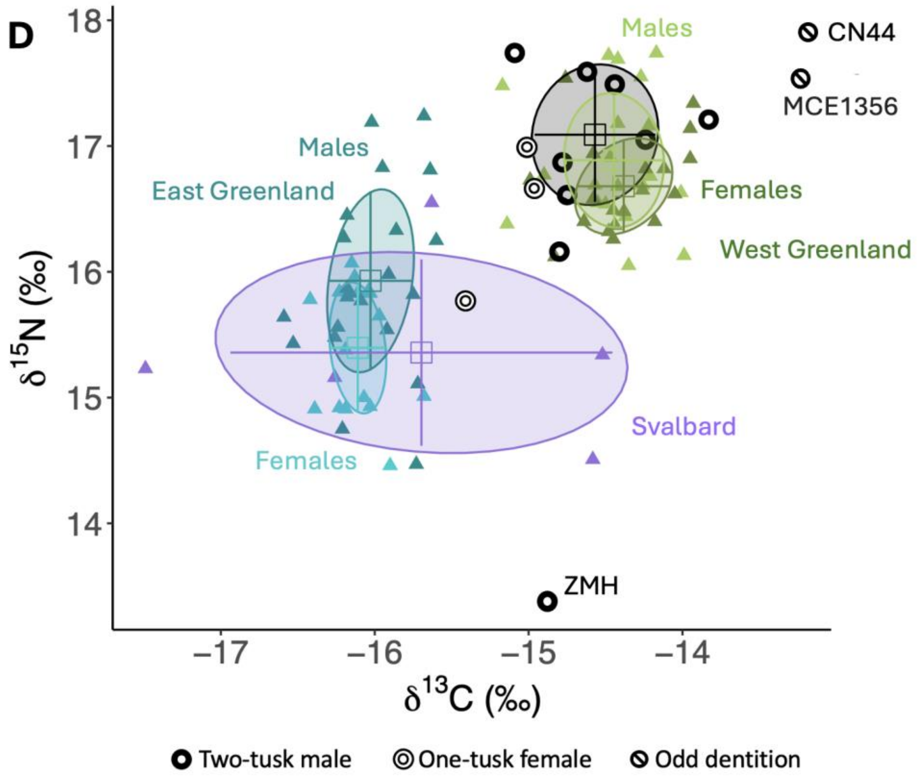


**Supplementary figure S2. Bone collagen $\delta^{13}$C and $\delta^{15}$N analysis with the reference data split by geography and sex**. The 14 anomalous-tusked individuals analysed are shown in black. The narwhal reference data are from: West Greenland (green): 20 females, 19 males; East Greenland (teal): 16 females, 22 males. The five Svalbard individuals (purple) were not genetically sexed. Shaded ovals indicate Bayesian standard ellipse areas for each group ($SEA_B$). Mean (square) and SD (error bars) are indicated. The three anomalous-tusked individuals discussed by name in the main text are indicated.

**Table S1. Sample and biomolecular data overview.** Included are the 15 anomalous-tusked and five Svalbard individuals for which novel data were generated for this study. Institution indicates where the specimen is housed: Natural History Museum of Denmark, University of Copenhagen (NHMD); Varde Museum; Greenland Institute of Natural Resources (GINR); Museum der Natur Hamburg of the Leibniz Institute for the Analysis of Biodiversity Change (LIB); Norwegian Polar Institute (NPI); Natural History Museum, University of Oslo (NHM); Swedish Museum of Natural History (NMR). TEAL (Trent Environmental Archaeology Lab) ID indicates the sample ID of University of Trent, where the stable isotope data were generated. Stable isotope (SI) data include $\delta^{13}$C corrected for the Suess effect. Geographic region refers to West Greenland (WG), east of Greenland (EG), or Greenland (GL, if specific region of origin is unknown).

| **Institution** | **Institution ID** | **TEAL ID** | **Sex** | **SI** | **$\delta^{13}$C (‰)** | **$\delta^{13}$C Suess (‰)** | **$\delta^{15}$N (‰)** | **C/N** | **Year for Suess correction** | **Collection year** | **Registration year** | **Locality and region** | **Comments** |
|---|---|---|---|---|---|---|---|---|---|---|---|---|---|
| NHMD | M08-CN1x | 5295 | M | new | -14.8 | -14.8 | 16.16 | 3.23 | 1800 | | 1800 | GL | specimen from the Royal Art Chamber, could be even older (as early as 1700s) |
| NHMD | M08-CN10x | 5296 | M | new | -14.62 | -14.58 | 17.59 | 3.46 | 1883 | 1883 | 1884 | Itivdliarsuk, WG | |
| NHMD | M08-CN11x | 5297 | M | new | -14.78 | -14.74 | 16.87 | 3.41 | 1884 | 1884 | 1884 | Upernavik, WG | |
| NHMD | M08-CN35x | 5298 | M | new | -14.49 | -14.44 | 17.49 | 3.56 | 1897 | | 1897 | GL | |
| NHMD | M08-CN58 | 5301 | M | new | -15.19 | -15.09 | 17.74 | 3.41 | 1921 | | 1921 | Kap York, WG | |
| NHMD | M08-CN76 | 5302 | M | new | -14.56 | -14.24 | 17.05 | 3.23 | 1965 | | 1965 | GL | bought from a person in Copenhagen in 1965 |
| NHMD | M08-CN2x | 10501 | M | new | -13.83 | -13.83 | 17.21 | 3.32 | 1800 | | 1800 | Omenakfjord, WG | specimen from the Royal Art Chamber, could be even older (as early as 1700s) |
| Varde Museerne | NA | 16393 | M | new | -14.75 | -14.75 | 16.61 | 3.4 | NA | | | Greenland | |

| | | | | | | | | | | | | | |
|---|---|---|---|---|---|---|---|---|---|---|---|---|---|
| LIB | ZMH-S-10192 | 23849 | M | new | -14.88 | -14.88 | 13.38 | 3.24 | 1684 | 1684 | | Greenland Sea (between Svalbard and EG) | SI run in duplicate with nearly same values found (δ13C: -14.92, δ15N: 13.28) |
| GINR | 1029 | NA | M | NA | NA | NA | NA | NA | 2009 | 2009 | | Scoresby Sound, EG | |
| GINR | 1196 | 17466 | F | new | -15.73 | -15.01 | 16.99 | 3.13 | 1996 | 1996 | | Kitsissuarsuit, WG | correction factor dentine to bone applied |
| GINR | 1197 | 17467 | F | new | -15.68 | -14.96 | 16.66 | 3.15 | 1996 | 1996 | | Kitsissuarsuit, WG | correction factor dentine to bone applied |
| NHMD/GINR | 938 | 11295 | F (Garde & Heide-Jørgensen, 2022) | (Rey-Iglesia et al., 2022) | -16.68 | -15.41 | 15.77 | 3.18 | 2017 | 2017 | | Scoresby Sound, EG | |
| NHMD | M08-CN44 | 604 | M (Vicari et al., 2022) | (Vicari et al., 2022) | -13.52 | -13.22 | 17.89 | 3.19 | 1963 | NA | 1963 | GL | |
| NHMD | MCE1356 | 575 | M (Skovrind et al., 2019) | (Skovrind et al., 2019) | -13.84 | -13.23 | 17.52 | 3.2 | 1986 | 1986 | | Kitsissuarsuit, WG | Also named Narluga |
| NPI | M.sv.4/SV08 | 17468 | NA | new | -16.26 | -16.26 | 15.16 | 3.1 | NA | | | Svalbard | sampled in 2012, also named CGG-1-012748 |
| NPI | MM2014/01 | 17469 | M | new | -16.8 | -15.63 | 16.55 | 3.14 | 2014 | 2014 | | Recherchefjorden, Svalbard | subadult male skeleton found onshore, also named CGG-1-017632 |
| NHMO | NHMO-DMA-46574/1-O | 14143 | NA | new | -17.54 | -17.49 | 15.23 | 3.53 | 1900 | | | Svalbard | uncertainty about date, also named CGG-1-024518 |

| | | | | | | | | | | | | | |
|---|---|---|---|---|---|---|---|---|---|---|---|---|---|
| NRM | NRM-MA558407 | 14545 | NA | new | -14.56 | -14.52 | 15.34 | 3.19 | 1886 | 1886 | | Svalbard | |
| Svalbard Museum | SVB 7773 | 17902 | NA | new | -14.58 | -14.58 | 14.51 | 3.14 | NA | | 2014 | Svalbard | fragment of tusk, correction factor dentine to bone applied |

**Table S1. Mapping statistics and genetic sexing.** The number of unique reads corresponds to the number of remaining mapped reads after taking out the PCR duplicates. Genetic sex was determined by estimating the X chromosome:autosome (A) coverage ratio (X:A ratio).

| **Specimen** | **Nb of mapped reads** | **Nb of unique reads** | **Mean coverage X** | **Mean coverage A** | **Mean X:A ratio** |
|---|---|---|---|---|---|
| M08-CN1x | 2882876 | 2524340 | 0.03 | 0.05 | 0.56 |
| M08-CN10x | 720006 | 620759 | 0.01 | 0.02 | 0.52 |
| M08-CN11x | 1091075 | 966276 | 0.02 | 0.03 | 0.54 |
| M08-CN35x | 2599245 | 2270864 | 0.03 | 0.05 | 0.55 |
| M08-CN58 | 4584063 | 4050866 | 0.06 | 0.1 | 0.55 |
| M08-CN76 | 2471885 | 2182721 | 0.04 | 0.07 | 0.55 |
| M08-CN2x | 4931349 | 4334837 | 0.06 | 0.12 | 0.53 |
| NA | 1057809 | 902396 | 0.02 | 0.04 | 0.5 |
| ZMH-S-10192 | 6061129 | 5924311 | 0.08 | 0.14 | 0.55 |
| 1029 | 76477141 | 60425812 | 2.05 | 3.9 | 0.53 |
| 1196 | 6265412 | 5770557 | 0.24 | 0.24 | 1.01 |

| | | | | | |
|---|---|---|---|---|---|
| 1197 | 4151302 | 3837687 | 0.13 | 0.12 | 1.07 |

### Supplementary text

#### Specimens

Additional information on three of the anomalous tusk specimens is provided here.

Two putatively female one-tusked narwhals from West Greenland were included in our analysis (table 1, supplementary table S1, supplementary figure S1b). The tusks were part of a batch of eight tusks - four identified as female by hunters - purchased in September 1996 by the general store (Kongelige Grønlandske Hansel) in Kitsissuarsuit, and brought to Nuuk. Incidentally, Kitsissuarsuit is the settlement where the Narluga, the first-generation male hybrid between a narwhal and a beluga, was collected (Heide-Jørgensen & Reeves, 1993). The chairman of the hunter’s organisation in Kitsissuarsuit was contacted by Aqqalu Rosing-Asvid for further information; he mentioned that several females with tusks had been caught that year, and that tusked females had been caught previously around Kitsisuarsuit (this is not the case in other areas). The tusk from specimen 1197 is relatively short (115 cm), but it has a clearly defined root (the part of the tusk that is embedded in the skull, which is smooth and without a defined root until the tusk stops growing in length, supplementary figure S1b). Normally, a 115 cm tusk is hollow in most of its length, but this tusk was filled out with dentine and was about ½ kg (40%) heavier than a normal tusk the same length. The tusk of specimen 1196 was longer (172 cm), and it also had a defined root, and was unusually slim. Another unusual feature was the reddish colour of the tusk, which may originate from some kind of algae pigment. Two of the other eight tusks were also relatively short (160 cm, 172 cm), but with a root, but they were not available for sampling.

The oldest of the sampled two-tusked individuals was a specimen caught in the Greenland Sea in 1684 (table 1, supplementary table S1). It was presumed to be a female due to its association with a foetus (figure 1). The text of a contemporaneous news bulletin explains the origin of the *unicorn*. It praises the rarity and high trade value of the two-tusk specimen: *It is so rare / that no one has seen such a fish / let alone caught one*. The caption of the depicted foetus (figure 1) leaves room for interpretation: *A young unicorn / as such found in the fish / the length and thickness match the original*. The text thus avoids to explicitly state that the foetus depicted was taken from the two-tusked narwhal.

### Methods and results

#### DNA

Laboratory work

We carried out DNA extractions following a silica column-based protocol with the binding buffer from (Allentoft et al., 2015). We incubated dentine powder overnight at 37°C with constant rotation in 1 mL of extraction buffer. After the overnight incubation, the supernatant was added to a 30 KDa Amicon® Ultra-4. The sample was spun down at 4,000 rpm, until the supernatant was concentrated down to 70 µl. The concentrate was combined with a 10x modified Qiagen PB buffer as described in (Allentoft et al., 2015) and purified using Monarch columns (NEB). After binding, we performed two washes with Qiagen PE. DNA elution was performed in two steps; for each step, we added 25 µl of Qiagen EB buffer to the Monarch column, incubated for 5 min at room temperature, and centrifuged at 13,000 rpm (max speed) for 1 min.

Prior to the library build, DNA extracts were treated with Thermolabile USER II enzyme (NEB). For each sample, the USER reaction was performed in 16 µl, with 2.4 µl of the Thermolabile USER II enzyme and 13.6 µl of each extract with an incubation time of 3 h at 37°C. USER treated DNA extracts were purified using Monarch columns (NEB).

DNA extracts were transformed into single stranded DNA (ssDNA) sequencing libraries as in (Kapp et al., 2021) and double-indexed using KAPA HiFi HotStart Uracil+ ReadyMix (KAPA Biosystems). The resulting indexed libraries were quantified on a Qubit™ dsDNA HS (Invitrogen) and quality checked in the Agilent Fragment Analyzer™. Indexed libraries were shotgun sequenced on an Illumina NovaSeq 6000 with the 150 base pairs (bp) PE technology.

For the two-tusked individual from which only soft tissue was available for analysis, we used the Qiagen DNA blood & tissue kit, using the guidelines from the manufacturer with minor modifications (increase of the Proteinase K volume to 40 µl, incubation time was extended to 24 h, and tissue was manually lysed with a pilar after the night of incubation). This DNA sample was sent to Novogene for library build and sequencing on an Illumina NovaSeq 6000 with the 150 bp PE technology.

### Genetic sexing

First, we identified scaffolds putatively originating from the sex chromosomes in the narwhal reference genome assembly (GCF_005190385.1). For this, we aligned the narwhal reference genome to the Cow X (Genbank accession: AC_000187.1) and Human Y (Genbank accession: NC_000024.10) sex chromosomes using satsuma synteny v.2.1 (Grabherr et al., 2010) with default parameters. Then we mapped the raw reads to the narwhal reference assembly using paleomix v.1.3.8 (Schubert et al., 2014) with BWA v.0.7.15 backtrack algorithm (Li & Durbin, 2009), requiring a minimum mapping quality of 30 and removing duplicates. We removed any risk of human contamination by mapping the narwhal fastq files to the human genome (hg38.UCSC.fasta) to generate bam files only keeping reads not mapping to the human genome. We then followed the SeXY pipeline (https://github.com/andreidae/SeXY) (Cabrera et al., 2022) and used the "Coverage_calculation.sh" script to estimate the average coverage of sequencing reads aligning to the X chromosome and the autosomes. We determined the sex of each specimen by estimating the X chromosome:autosome coverage ratio (X:A ratio). Specimens with X:A ratio < 0.7 were determined as males, and those with X:A ratio > 0.8 as females. Mapping statistics, mean ratios and coverage can be found in Table S2.

### Stable isotopes

#### Laboratory work

Powdered samples underwent lipid extraction with 3 repetitions of 10 mL 2:1 chloroform/methanol (*v*/*v*) under sonication for 1 h. After removing the solvent, we dried the samples under normal atmospheric pressure for 24 h. We demineralized the samples in 10 mL of 0.5 M HCl for 1 h while agitating them by an orbital shaker. After demineralization, we rinsed the samples to neutrality with Type I water, and heated them at 75 °C for 36 h in 0.01 M HCl to solubilize the collagen. The water soluble collagen was transferred to a 4 mL glass vial, frozen, and lyophilized. Upon analysis there was an observed correlation between $\delta^{13}C$ and $C:N_{atomic}$, indicating the presence of residual lipids. To extract these lipids we solubilized the lyophilized collagen, and add chloroform and methanol to obtain a 10 mL solution of 10:5:4 chloroform/methanol/water (*v*/*v*/*v*) under sonication for 1 h. After centrifugation, the layer containing solubilized collagen and some methanol was collected and transferred to a new 4 mL glass vial; any residual methanol was evaporated from the solution at 60 °C for 24 h. We freeze dried and weighed the samples before adding 0.5 mg into tin capsules for elemental and isotopic analysis. We determined carbon and nitrogen stable isotopic and elemental compositions using a Euro EA 3000 Elemental Analyzer (Euro Vector SpA) coupled to a Nu Horizon (Nu Instruments, UK) continuous flow isotope ratio mass spectrometer at the Water Quality Centre at Trent University, Canada. We calibrated stable carbon and nitrogen isotopic compositions relative to the VPDB and AIR scales using USGS40 and USGS41a (glutamic acid) (Qi et al., 2003, 2016). Analytical uncertainty was monitored using four internal reference materials in addition to USGS40 and USGS41a: SRM-1 (caribou bone collagen, long-term average $\delta^{13}C = -19.40 \pm 0.08$ ‰, $\delta^{15}N = +1.82 \pm 0.11$ ‰), SRM-2 (walrus bone collagen, long-term average $\delta^{13}C = -14.82 \pm 0.06$ ‰, $\delta^{15}N = +15.59 \pm 0.14$ ‰), SRM-14 (polar bear bone collagen, long-term average $\delta^{13}C = -13.68 \pm 0.08$ ‰, $\delta^{15}N = +21.61 \pm 0.16$ ‰), and SRM-17 (phenylalanine, long-term average $\delta^{13}C = -12.45 \pm 0.04$ ‰, $\delta^{15}N = +3.17 \pm 0.15$ ‰). For samples analyzed in duplicate, the mean difference between pairs was 0.06 ‰ for $\delta^{13}C$ and 0.11 ‰ for $\delta^{15}N$. Standard uncertainty was determined to be ±0.11 ‰ for $\delta^{13}C$ and

±0.21 ‰ for $\delta^{15}$N. We also adjusted the $\delta^{13}$C values to correct for the change in atmospheric and oceanic dissolved inorganic carbon that has occurred since the late 19th century due to industrialization (the "Suess Effect"; (Keeling et al., 1979; Quay et al., 1992)), following (Szpak et al., 2020),, although we used 0.014 for the annual rate at which $\delta^{13}$C has declined for a particular water body (Mellon, 2018). The formula used is as follows:

$$\delta 13C\ Suess = 0.014\ x\ (e(collection\ year - 1850)^{0.027}) +\ \delta 13C$$

We did not correct $\delta^{13}$C values for the Suess effect for specimens for which the exact collection date is unknown but originating from the 1800s, according to the museum's records, because this phenomenon had no effect prior to around 1850. We used the registration date for specimens for which collection date was not available. For atmospheric CO2 the shift between 1850 and 1930 is 0.3 ‰ (more likely 0.1 ‰ in the ocean), and thus a collection date earlier than the entry date would have little effect on our results.

To analyse bone and dentine (tusk) data together, dentine isotopic values need to be translated into bone-equivalents using a correction factor, and we thus applied a correction factor to the $\delta^{13}$C and $\delta^{15}$N values (0.81 and 0.49 were subtracted respectively) obtained from the two tusks (1196 and 1197) (Rey-Iglesia et al., 2022).

Sex-based isotopic differentiation

We tested for dietary differences between the two-tusked individuals (n=8) and reference males and females from West Greenland separately ($n_M$=19, $n_F$=20), by comparing their $\delta^{13}$C and $\delta^{15}$N using Student's t-tests. Note that isotopic differentiation has been observed between males and females in East Greenland, where males have higher $\delta^{15}$N than females (Louis et al., 2021), but not in West Greenland. However, we wanted to test whether there were any differences with the two-tusked individuals, and also visually compare the one-tusked females and individuals with highly unusual dentition with the reference panel of their respective regions.

We did not find any significant difference between the two-tusked narwhals and the West Greenland reference data when split by sex: males ($t = 1.20$, $df = 10.84$, $P = 0.26$ for $\delta^{13}C$; $t = -1.98$, $df = 10.02$, $P = 0.08$ for $\delta^{15}N$), or females ($t = 0.80$, $df = 10.86$, $P = 0.44$ for $\delta^{13}C$; $t = -0.91$, $df = 12.73$, $P = 0.38$ for $\delta^{15}N$, supplementary figure S2). The isotopic niche area of the West Greenland two-tusked narwhals was larger, but not significantly different, than those of West Greenland narwhal males ($p=0.95$, $SEA_B=0.63$ ‰$^2$) and females ($p=0.73$,$SEA_B=0.55$ ‰$^2$, supplementary figure S2). The ellipse overlap of the two-tusked narwhals and the reference material for West Greenland males was 20%, and the ellipse overlap between the two-tusked narwhals and the females was 49%.
